# Longitudinal comparison of the developing gut virome in infants and their mothers

**DOI:** 10.1101/2022.05.13.491764

**Authors:** Andrea C Granados, Catherine Ley, William A. Walters, Scot Federman, Yale Santos, Thomas Haggerty, Alicia Sotomayor-Gonzalez, Venice Servellita, Ruth E Ley, Julie Parsonnet, Charles Y Chiu

**Author notes:** co-senior authors (Julie Parsonnet, Charles Y Chiu). Corresponding author: Charles Y Chiu MD PhD, University of California San Francisco, 185 Berry St, Box 0134, San Francisco, CA, 94158.

## Abstract

The virome of the human gut and its development in early life are poorly understood. Here we performed viral metagenomic sequencing on stool samples from a multiethnic, socioeconomically diverse cohort of 53 infants collected longitudinally over their first 3 years of life and their mothers to investigate and compare their viromes. The asymptomatic infant virome consisted of bacteriophages, dietary/environmental viruses, and human pathogenic viruses, in contrast to the material virome, in which sequence reads from human pathogenic viruses were absent or present at extremely low levels. Picornaviruses and phages in the family *Microviridae* (microviruses) dominated the infant virome, while microviruses and tomato mosaic virus dominated the maternal virome. As the infants aged, the human pathogenic and dietary/environmental virus components remained distinct from the materal virome, while the phage component evolved to become more similar. However, the composition of the evolving infant virome was not determined by the mother and was still maturing to the adult virome at three years of age.

**Importance:** The development of the human gut virome in early childhood is poorly understood. Here we use viral metagenomic sequencing in a cohort of 53 infants to the characterize their gut viromes and compare them to their mothers’.. This study finds that the infant virome consists of phages and human pathogenic viruses in asymptomatic individuals and is still maturing into the adult virome at three years of age.

## Introduction

Within hours of delivery, infants are colonized by bacteria and viruses (Lim et al., 2015; Maqsood et al., 2019; Liang et al., 2020). Unlike the bacterial microbiome, little is known about the gastrointestinal virome (bacteriophages, archaeal, and eukaryotic viruses) and its influence on growth and development in early childhood. Several technical limitations, as described below, have hampered this field. Selective enrichment of viral sequences is necessary to avoid preferentially sequencing bacterial or host DNA. In prior metagenomic analyses, a large proportion of viral sequences have not agreed to known reference genomes (Liang & Bushman, 2021), perhaps because sequences from bacteriophages (phages), which typically predominate the virome, are poorly represented in existing databases such as National Center for Biotechnology Information (NCBI)’s GenBank. RNA viruses in particular have been poorly studied because RNA is less stable in samples collected primarily for DNA-based metagenomic studies, and DNA contamination from laboratory reagents can also interfere with virome analysis (Santiago-Rodriguez & Hollister, 2019). Consequently, RNA viruses remain under sampled in most virome studies (Krishnamurthy et al., 2016; Callanan et al., 2020). To combat these problems, we and others have developed targeted viral enrichment approaches and improved bioinformatics methods for annotating sequences and analyzing metagenomic data (Legoff et al, 2017; Miller, et al., 2019; Santiago-Rodriguez & Hollister, 2019).

Most studies in infants that investigate the early development of the virome have included small sample sizes over short time frames (a few months to a year) (Lim et al., 2015; Masqood et al., 2019, Taboada et al., 2021). These studies have identified the appearance of phages such as *Siphoviridae, Inoviridae, Myoviridae, Podoviridae* and *Microviridae*, as well as the sporadic occurrence of eukaryotic viruses such as *Adenoviridae, Astroviridae, Caliciviridae, Parvoviridae, Picornaviridae*, and *Polyomaviridae* (Rascovan et al., 2016; Santiago-Rodriguez & Hollister, 2019; Shokoporov et al., 2019; Zuo et al., 2020; Lim et al., 2015). Longer longitudinal investigations of the virome in healthy and diverse populations and comparative studies in children and adults are needed to expand our knowledge of human-microbe interactions in childhood development. In this study, we utilized metagenomic viral sequencing to characterize the gastrointestinal virome in longitudinally collected stools from an ethnically and socioeconomically diverse prospective cohort of infants in California over the first three years of life. The availability of this cohort offers a unique opportunity to study the longitudinal progression of the infant virome in early childhood and to compare the viromes of mother-infant pairs.

## Methods

### Ethics and clinical and epidemiological metadata collection

Stanford’s Outcomes Research in Kids (STORK) is a multiethnic cohort of mothers and their babies from the second trimester of pregnancy through their third birthday. As previously described (Ley et al., 2016), subject recruitment for the cohort was performed at two obstetric clinics, one at the Stanford School of Medicine and the other within Santa Clara Valley Medical Center, via posters and flyers. Any healthy pregnant woman aged 18-42 years with a single fetus who wished to participate was enrolled in the study; babies were enrolled approximately two weeks post-delivery, before the initial household visit. A randomized intervention of triclosan and triclocarban-containing (TC) household and personal cleaning products over the first 12 months of life was nested within the cohort (Ley et al., 2018).

Relative to the overall population of the United States, the cohort is overrepresented by Hispanic/LatinX families and families of lower socioeconomic status. Households were visited every four months (M1-2; B1-9). At the initial household visit (second trimester, M1), the mothers were given a detailed household questionnaire including demographic information, information on the household structure (including inhabitants), and indicators of socioeconomic status. Starting at the first household visit after delivery (B1), stool, urine, and blood samples were obtained from both the mother and the baby, placed on ice packs, and frozen within 24 h of collection (Figure 1A). Frozen stool samples were stored at −80 °C before processing. Weekly automated telephone or email surveys were used to assess for signs of infectious disease in the children. These surveys recorded total days per week of vomiting, fever, diarrhea, and symptoms suggesting upper respiratory infection (URI), including cough, nasal congestion, and ear pulling. If the children were reported as healthy, parents were asked about milk intake and amount of sleep to ensure equal amounts of time were spent answering questions regardless of the child’s health status. A medical chart review was performed every six months as available.

**Figure 1.**
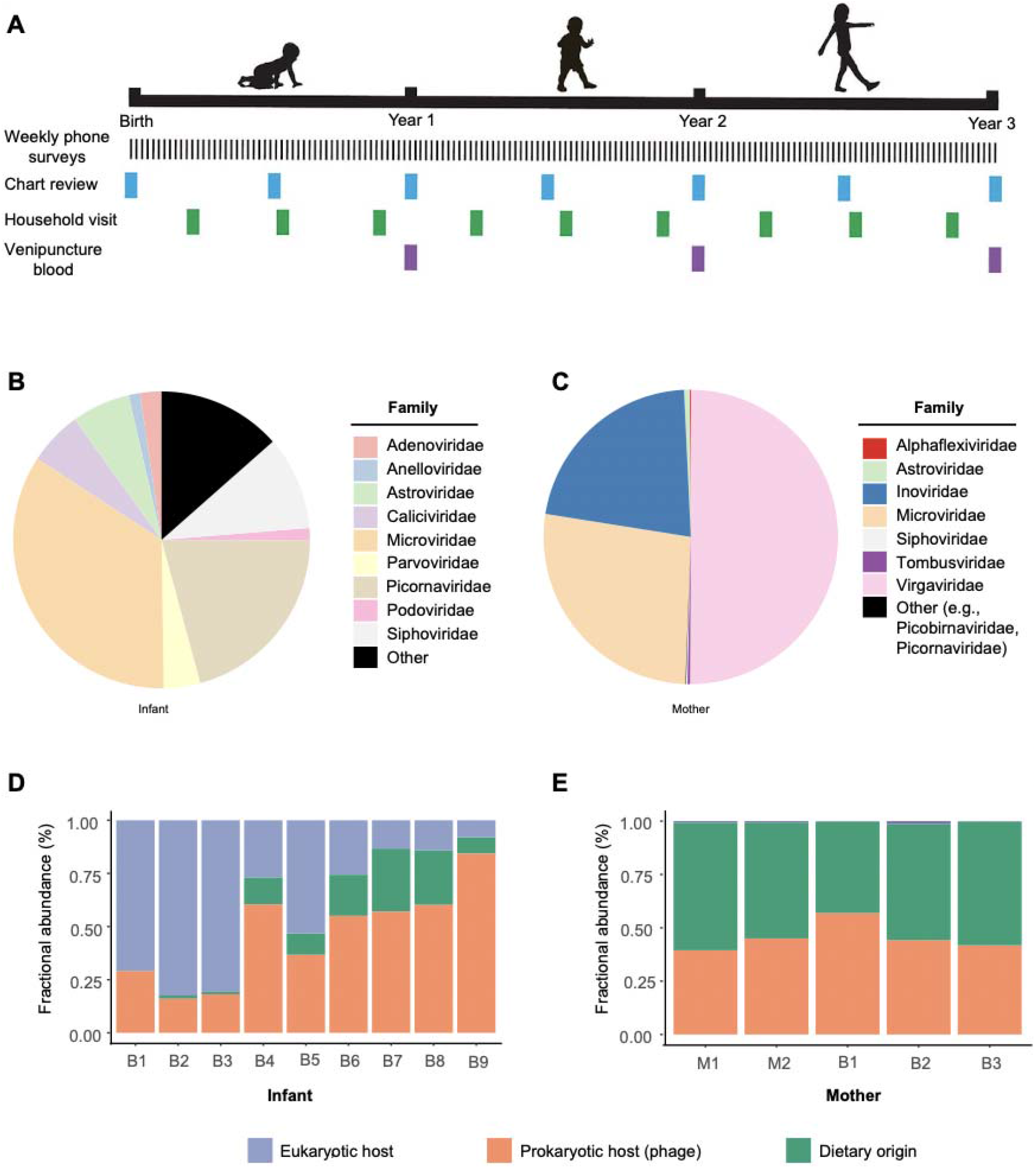
Overview of STORK sample and data collection and virome composition in infants and mothers over time. **(A)** Stool samples for mothers and infants were collected at each household visit. **(B)** Distribution of the most abundant viral families in infants. ‘Other’ consists of *Papillomaviridae, Polyomaviridae, Hepadnaviridae, Herpesviridae, Reoviridae, Retroviridae, Circoviridae, Astroviridae*, and *Picobirnaviridae*. **(C)** Distribution of the most abundant viral families in mothers. ‘Other’ consists of *Papillomaviridae, Polyomaviridae, Hepadnaviridae, Herpesviridae, Reoviridae, Retroviridae, Circoviridae, Astroviridae*, and *Picobirnaviridae*. **(D,E)** Fractional abundance of eukaryotic (human pathogenic), diet/environmental, and prokaryotic viruses in infants and mothers.

Recruitment of subjects, documentation of informed consent, collection of stool samples, sample processing, and metagenomic analyses were carried out with Institutional Review Board approval from Stanford University (Stanford, CA, USA), the Santa Clara Valley Medical Center (SCVMC) (San Jose, CA, USA) and University of California, San Francisco (San Francisco, CA, USA).

### Extraction

Stool samples were diluted 20% in phosphate-buffered saline (PBS) (1500 ml) and centrifuged for 5 min at 10,000g, followed by filtration using a 0.45 uM filter and treatment utilizing a nuclease cocktail of TURBO DNase (Invitrogen), Baseline Zero DNase (Ambion), Benzonase (Novagen) and RNase A (Roche) for 30 min at 37°C. This procedure digests host cell and non-protected (naked) viral nucleic acids while maintaining viral nucleic acids in particles protected from nucleases’ action. Nuclease activity was immediately inactivated by adding guanidium-thiocyanate containing lysis buffer (Qiagen), followed by total nucleic acid extraction of 400 ul of pretreated stool using the EZ1 Virus Mini Kit v2.0 (Qiagen). Extracts were eluted in 60 ul volume.

### Library Preparation

Amplified cDNA was prepared using random nonamer primers attached to a primer linker sequence with 25 cycles of PCR amplification, as previously described (Legoff et al, 2017). Amplified cDNA was purified using AMPure XP beads (Beckman-Coulter) on the EpMotion 5075 (Eppendorf). Purified dsDNA (2 ng) was used for NGS library generation using the Nextera Flex kit (Illumina) and purified using AMPure XP beads (Beckman Coulter) on the EpMotion 5073 (Eppendorf). Libraries were quantified using the Qubit dsDNA HS Assay (Thermofisher Scientific) on the Qubit Flex (Thermofisher Scientific). Samples were sequenced on the NovaSeq 6000 (Illumina) using 150bp paired-end sequencing at the UCSF Center for Advanced Technology (CAT). Samples were batched (80-96 per lane), with negative control (nuclease-free water) and a reference standard containing six non-pathogenic microorganisms.

### Metagenomic Analysis

Metagenomic next-generation sequencing (mNGS) data from all samples were analyzed for viral nucleic acids using SURPI+ (v1.0.7-build.4) (Naccache et al., 2014; Miller, et al., 2019), a bioinformatics pipeline for pathogen detection and discovery from metagenomic data, modified to incorporate enhanced filtering and classification algorithms. The SNAP nucleotide aligner (Zaharia et al., 2011) was run using an edit distance of 16 against the National Center for Biotechnology Information (NCBI) nucleotide (NT) database filtered to contain the viral, bacterial, fungal, and parasitic reads of GenBank (March 2019), enabling the detection of reads with ≥90% identity to reference sequences in the database. The pre-established criterion for viral detection by SNAP was the presence of reads mapping to at least three non-overlapping regions of the viral genome. Jaccard presence/absence distances were calculated on viral/phage counts and imported into QIIME2 (2020.2 release; Bolyen et al., 2019) to create visualization artifacts with biplots showing the top 5 taxa driving clustering. Note that the values were not rarefied, as they were already fractional values normalized against background host/microbial sequence counts.

### Statistical Analysis

Alpha diversity metrics, including the Abundance and Shannon Diversity Index, were calculated using the R package vegan2.5-3 (Oksanen et al., 2017). Comparisons of virome or microbiome abundance, richness, and alpha diversity between groups were analyzed using the Kruskal-Wallis test, followed by the Dunn test for post hoc analyses.

## Results

The study included 53 babies born between August 2011 and July 2015 (Table 1). A majority (58.5%) of mothers were primarily Hispanic/LatinX, with more than a quarter (26.4%) not completing high school. The median household size was four people, with approximately 25% having no other children in the household. Of the 53 infants, 45% were female, and 98.1% (52/53) identified with a similar race and ethnicity as the mother. Two-thirds of babies were delivered vaginally; all but one had received at least some breast milk. Almost all babies (48/53; 90.6%) had at least one identified medical care visit for illness, and 60% of these were prescribed antibiotics at least once.

Babies provided a median of 9 stool samples over three years, starting two to three months after birth and every four months after that until their 3rd birthday (Table 2). Among those with medical care visits for illness, the mean time between the visit date and the next stool sampling date was 59 days. Mothers provided a median of 5 stool samples over a year, starting in the third trimester (1-2 samples), and a paired sample alongside the infants in the first year.

Viral metagenomic sequencing analysis of 454 stool samples from the infants yielded a total of 80.7 billion sequence reads, with a mean of 6.2 billion reads/sample (+/- 3.3 billion reads/sample, Table S1). Metagenomic sequence data were analyzed using the SURPI+ (sequence based ultra-rapid pathogen identification) bioinformatics pipeline (Miller, et al., 2019). The mean percentages of mapped viral, bacterial, and human preprocessed reads across all samples were 6.3%, 23.5%, and 13.7%, respectively (Table S1). An average of 33.0% of reads did not map to any reference sequence in NCBI NT. Analysis of 233 stool samples belonging to study participants’ mothers yielded a total of 9.8 billion sequence reads, with an average of 2.5 billion reads/sample (+/- 2.3 billion reads/sample, Table S2). The mean percentages of matched viral, bacterial, and human reads across all samples were 7.0%, 45.0%, and 10.0% of preprocessed reads, respectively (Table S2). An average of 37.1% of reads did not map to any reference sequence in NCBI NT.

Overall, we observed that the infant virome consists of various eukaryotic viruses, nearly all with a presumptive human host, and prokaryotic viruses (phages). The most common viruses were phages in the *Microviridae* (34.4%), *Picornaviridae* (20.7%), and *Siphoviridae* (10.2%) families (Figure 1B). In contrast, the mothers’ viromes consist predominantly of dietary-associated viruses (e.g., tomato mosaic virus), environmental viruses, and phages, the last group including *Virgaviridae* (50.0%), *Microviridae* (26.9%), and *Inoviridae* (21.8%) (Figure 1C). The infant virome shifted dynamically in composition from being eukaryotic virus dominant in the first year of life to phage dominant in the third year (Figure 1D). In contrast, the mothers’ viromes remained static before (M1, M2) and after birth (B1-B3) and were consisted of comparable proportions of dietary/environmental viruses and phages.

Based on an alpha diversity metric, we observed that the abundance and diversity of phages in the infants increased significantly between the first 1.5 years (B1-B4) and the third year (B7-B9) (p<0.05, Figure 2A-B). For eukaryotic viruses, however, we observed overall decreases in abundance and diversity from year one (B1-3) to years 2 and 3 (B5-9) (p<0.05 Figure 2C-D). Using principal coordinates analysis, we observed *Lactococcus* and gokushovirus phages driving the ordination of the younger and older infant prokaryotic viromes, respectively (Figure 2E). The eukaryotic virome ordination was consistently driven by picornaviruses (enterovirus A and B and parechovirus A) throughout (Figure 2F). We did not observe any change in the abundance of diet/environmental viruses as the infants aged (Figure S1A); however, diversity did significantly increase in the third year (B7-B9) compared to the first two time points (B1 and B2, p<0.0001) (Figure S1B). Although there was no definitive pattern with age in beta diversity, we did observe pepper mild mottled virus, tomato mosaic virus, tropical soda apple virus, chicken anemia virus, and chicken stool *Gemycircular* virus as diet-related viruses driving the ordination (Figure S1C).

**Figure 2.**
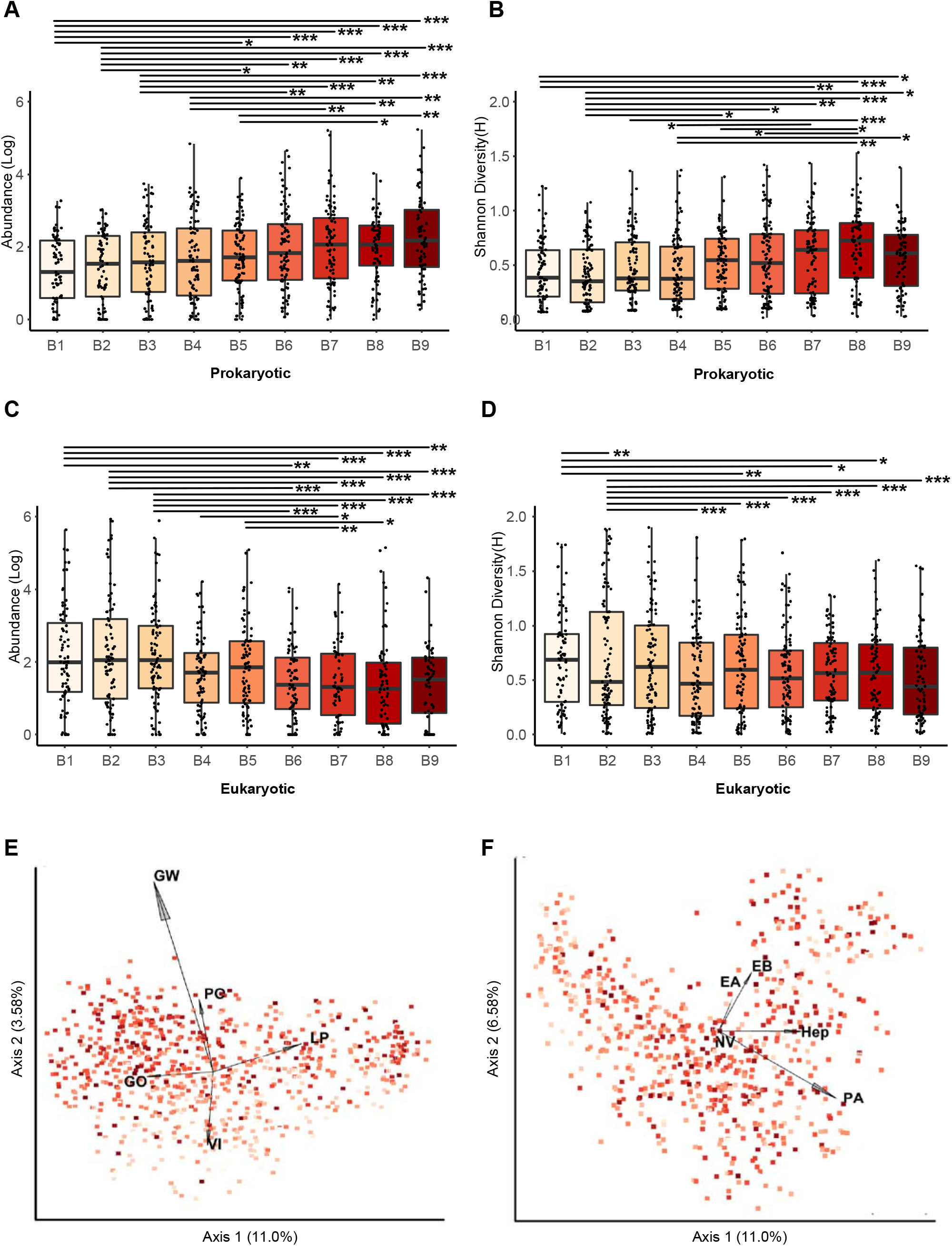
Normalized abundance, alpha diversity, and beta diversity over the first three years of life. Box and whiskers plot of **(A)** prokaryotic virus normalized abundance, **(B)** prokaryotic virus Shannon diversity, **(C)** eukaryotic virus normalized abundance, and **(D)** eukaryotic virus Shannon diversity. **(E)** Principal component analysis (PCA) plot showing infants clustered by Jaccard distances from prokaryotic viruses. **(F)** PCA plot of infants clustered by Jaccard distances of eukaryotic viruses. Samples are colorized with an age gradient (light to dark red representing age in days of the infants, range of 22 to 1166). The top 5 taxa driving clustering are shown as a biplot. Abbreviations: Hep, non-A-non-B hepatitis virus (2 uncategorized viral taxa in this group), PA: parechovirus A, NV: Norwalk virus, EA: enterovirus A, EB: enterovirus B. GW: Gokushovirus WZ-2015a, GO: Human gut gokushovirus, PO: Poophage MBI-2016a, LP: Lactococcus phage 936 sensu lato, VI: Vibrio phage JSF5. *, p<0.01; **, p<0.001; *** p<0.0001.

We observed that the infant virome over the study period primarily consisted of *Siphoviridae, Microviridae, Picornaviridae*, and *Anelloviridae* (Figure 3A-B), with decreasing amounts of *Myoviridae* and *Podoviridae* over time (Figure 3A-B). The top viral species identified in infants were human gut gokushoviruses, adenoviruses, noroviruses, parechoviruses, pepper mild mottled virus, *poophage* MBI2016a, and tropical soda apple mosaic virus (Figure 4). In general, the proportion of these most common species remained consistent over the years, except early in year 3 (B7), where we found an overrepresentation of parechovirus A (Figure 4).

**Figure 3.**
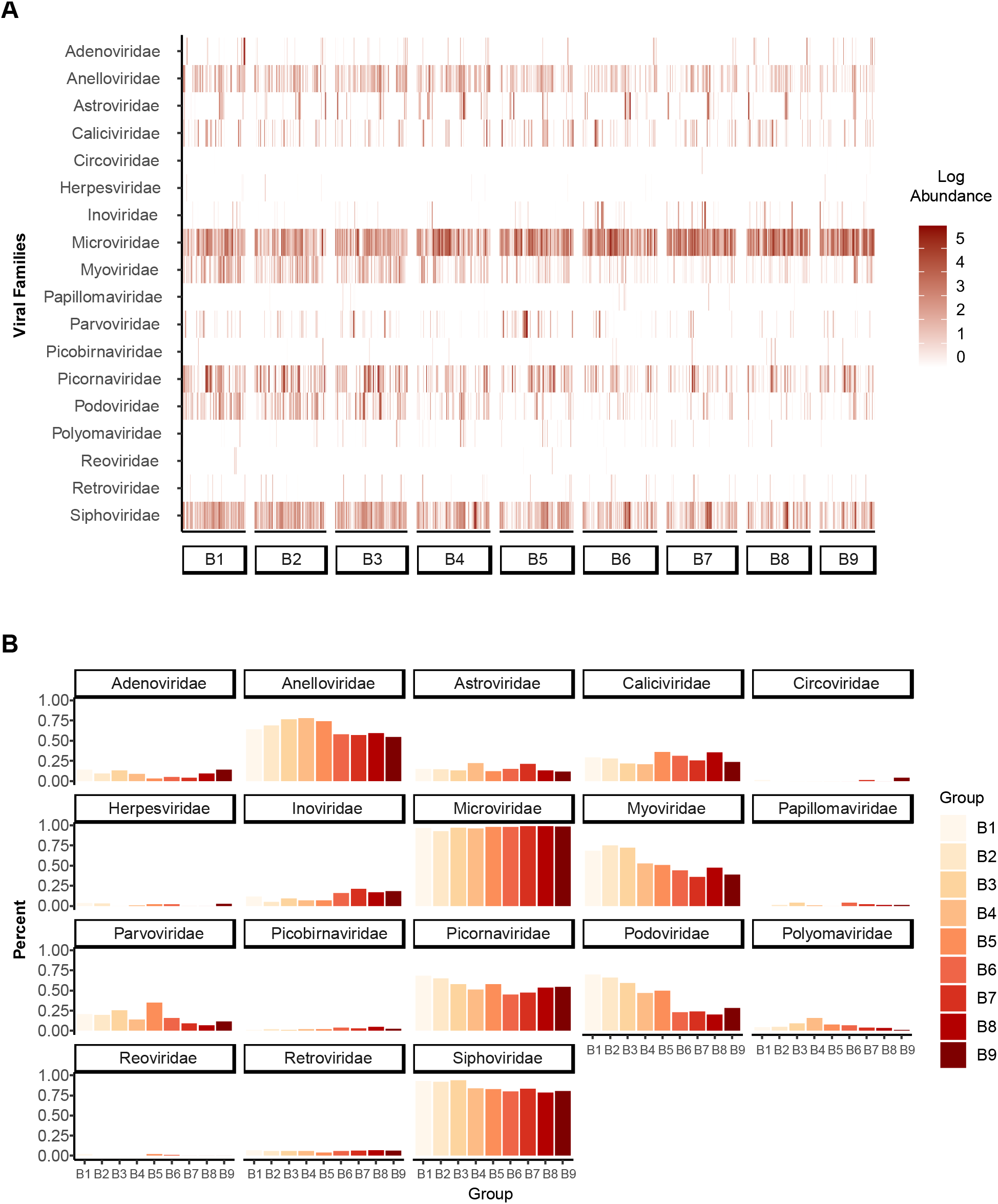
Distribution of viral families and top 12 viral species in infants. **(A)** Heatmap distribution of viral families in infants over time. **(B)** Percent relative abundance of viral families in infants over time.

**Figure 4.**
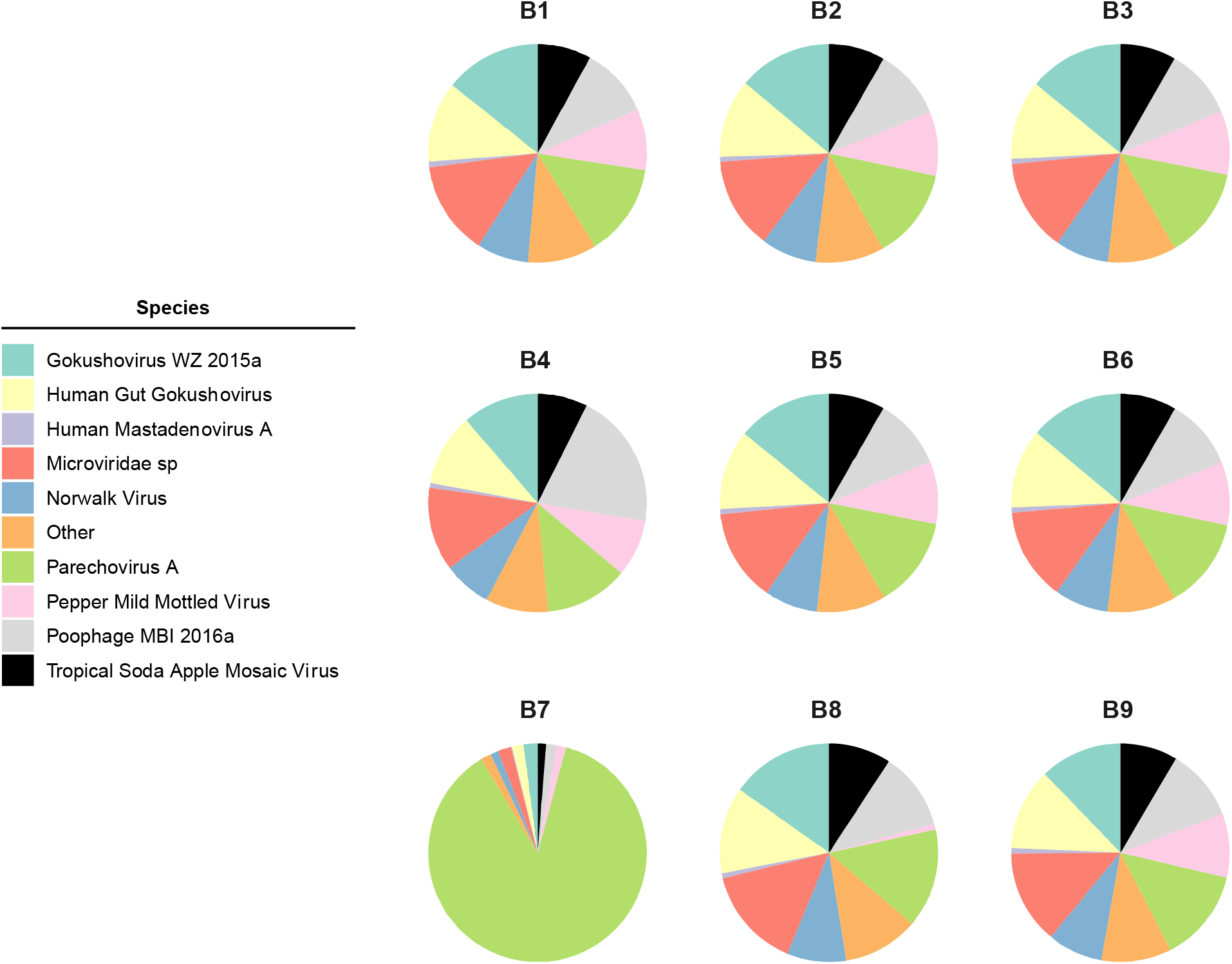
Composition of the top 10 viral species in infants.

We compared the alpha and beta diversity of the infants and the mothers in the first year of life (B1-B3). Phages were significantly more abundant in the maternal virome than the infant virome at all three time points (p<0.0001, Figure 5A); however, there was no difference in overall diversity in either virome population at any time point (p>0.05, Figure 5B). In contrast, the infant virome was more abundant in eukaryotic viruses than the maternal virome at all three time points (p<0.0001, Figure 5C), although eukaryotic diversity increased in the infant virome at the last time point (B3, Figure 5D, p=0.01).

**Figure 5.**
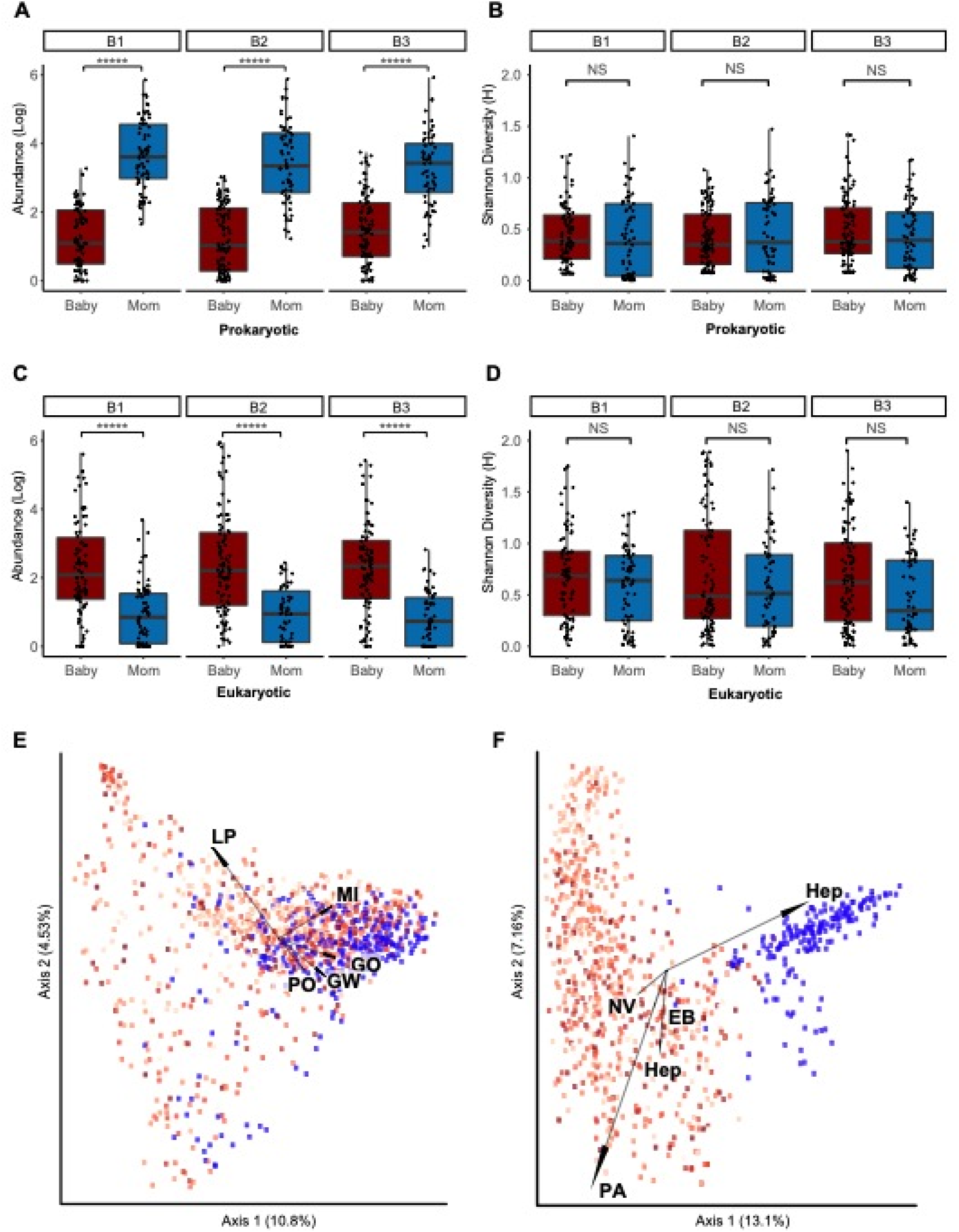
Comparison of alpha and beta diversity of infants and mothers in the first year. Box and whiskers plot of **(A)** prokaryotic virus abundance, **(B)** prokaryotic virus Shannon diversity, **(C)** eukaryotic virus abundance, and **(D)** eukaryotic virus Shannon diversity. **(E)** PCA plot of infant and mother samples, clustered by Jaccard distances of bacteriophage and archaeal viral counts. **(F)** PCA plot of infant and mother samples using human virus counts clustered with Jaccard distances. Samples are colorized with an age gradient (light to dark red representing age in days of the infants, range of 22 to 1166), and mothers are coloured in blue. The top 5 taxa driving clustering are shown as a biplot. Abbreviations: Hep, non-A-non-B hepatitis virus (2 uncategorized viral taxa in this group), PA: parechovirus A, NV: Norwalk virus, EA: enterovirus A, EB: enterovirus B. GW: Gokushovirus WZ-2015a, GO: Human gut gokushovirus, PO: Poophage MBI-2016a, LP: Lactococcus phage 936 sensu lato, VI: Vibrio phage JSF5. *, p<0.01; **, p<0.001; *** p<0.0001.

When observing the beta diversity of the phage virome in infants compared to the mothers, the infant phage population began to cluster alongside the maternal phage population with age (Figure 5E). The separation of the younger virome (B1) was affected by the presence of the *Lactococcus* phage. In contrast, the clustering of the older infants (B2-B3) and maternal phages was affected by the presence of Gokushoviruses, poophages, and other microvirus species (Figure 4E). Unlike phages, there was a clear separation of the eukaryotic virome between the infants and mothers (Figure 5F). The infant eukaryotic virome was separated from the maternal eukaryotic virome by the presence of noroviruses and picornaviruses (Enterovirus B, Parechovirus A) (Figure 5F). Alpha diversity of diet/environmental viruses in the mothers remained stable during and after pregnancy (p>0.05, Figure S2A, S2B). This component is consistently clustered separately from the infants, with the mothers having more tomato mosaic virus (Figure S2C).

Finally, we compared the infant virome at the third year (B9) to the mother’s last sequenced time point (B3). The maternal virome (phage and eukaryotic) was more abundant than the infants’ (p<0.0001, Figure 6A, 6C), however, the infant phages were more diverse than the mothers’ (p=0.0004, Figure 6B). There was no difference in the diversity of the eukaryotic virome in either population. Mothers harboured significantly fewer anelloviruses and picornaviruses than infants but more picobirnaviruses and chronic viruses like retroviruses and herpesviruses, while both populations contained a high abundance of microviruses (Figure 6E). Thus, the maturing infant virome was not fully developed by the third year of life. Additionally, we compared the beta diversity of infants to their mothers at the first (B1) and last (B9) timepoints to investigate whether longitudinally acquired diversity is driven by the maternal virome.Infant diversity was not determined by prior exposure to the mother **(Figure S3).**

**Figure 6.**
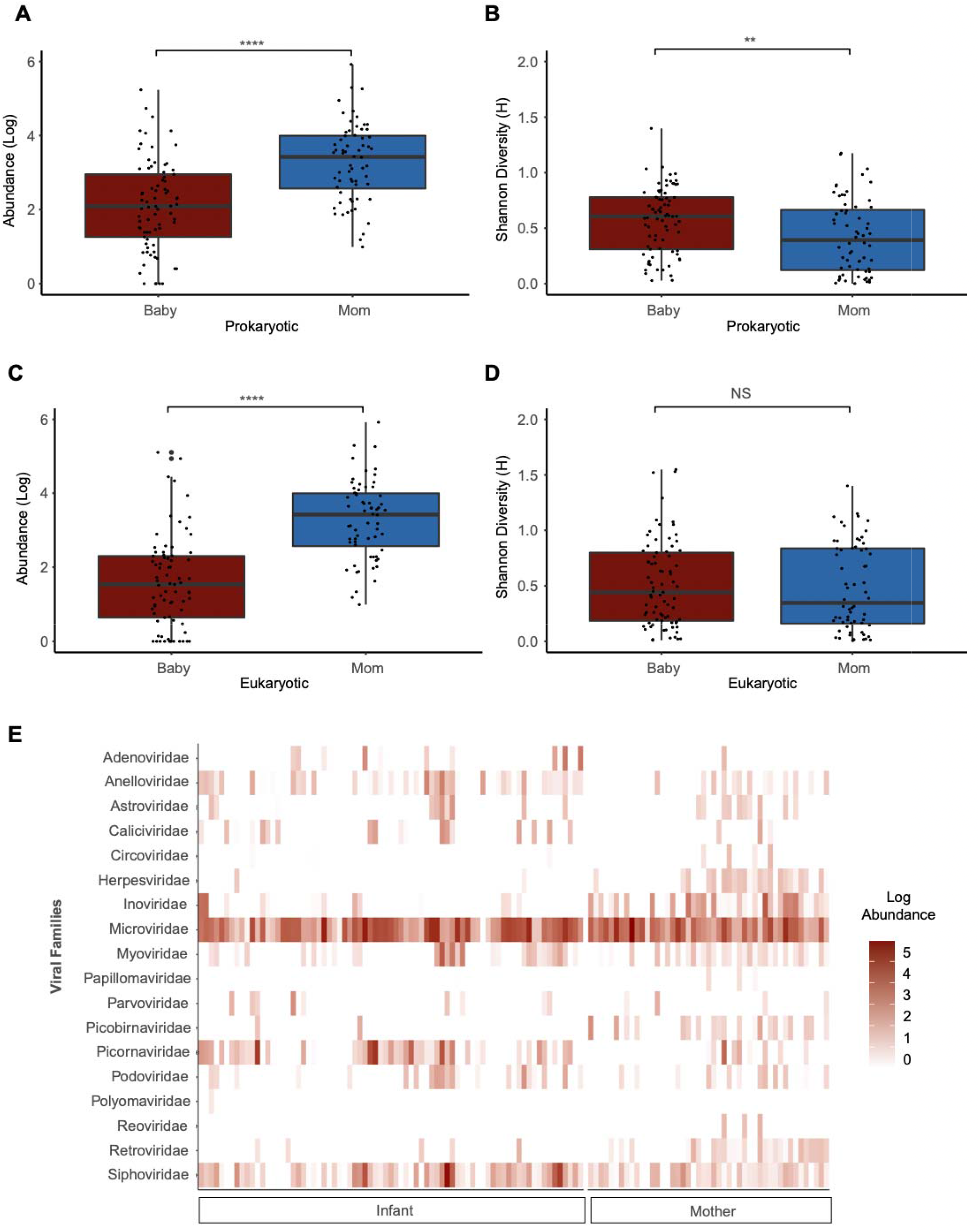
Comparison of mothers (B3) and infants (B9) sampled at the last time point. Box and whiskers plot of **(A)** prokaryotic virus abundance, **(B)** prokaryotic virus Shannon diversity, **(C)** eukaryotic virus abundance, and **(D)** eukaryotic virus Shannon diversity. **(E)** Heatmap of viral family abundances at B9 for infants and B3 for mothers. Abbreviations: ***, p<0.0001; ****, p<0.00001.

## Discussion

Here we use metagenomic sequencing of fecal viromes derived from cohort of 53 infants sampled quarterly over three years (July 2011 to August 2015) to characterize the development of the infant gut virome. We also described the gut viromes of their mothers in the infants’ first year to assess the progression of the adult virome. Unlike maternal virome, primarily composed of phage and dietary/environmental viruses, the infants’ virome bore a mixture of phage and eukaryotic viruses. As infants aged, the phage component of the virome shifted from being dominated by *Lactococcus* phage to Gokushoviruses, poophages and microvirus species approaching the composition of the maternal phage virome. In contrast, the eukaryotic (human-host) component of the infant virome was not mature and was still developing at year three.

The presence of *Lactococcus* phage drove the separation of the phage component of the infant virome from the maternal virome early in the child’s life (B1); these belong to the *Siphoviridae* family and infect the bacterium *Lactococcus lactis*, which are vital bacteria for the fermentation of milk/dairy products (Mahony et al., 2017). Our cohort reported breastfeeding for at least some time in 98.1% of infants, with 88.7% breastfeeding for at least three weeks, demonstrating the association between diet and virome diversity. As the children aged, *Microviridae--primarily* Gokoshuviruses and poophages-became the dominant phages in the gut (Liang et al., 2020; Lim et al., 2020; Takahashi et al., 2000). While little is known about the recently identified poophages (Santiago-Rodriguez et al., 2020), the Gokushoviruses have been recently described (Creasy et al., 2018; Székely and Breitbart, 2016; Kirchberger & Ochman, 2020). These phages were initially only identified in *Bdellovibrio* and *Chlamydophila* bacteria but have now been identified in *Escherichia coli*, surviving in the cytoplasm (Kirchberger & Ochman, 2020). The variety of lifestyle cycles permits these phages to persist in many environments, and they are most likely acquired by the infants from various sources (i.e., diet, environment). Future analyses will be required to determine how an increase in abundance of these microviruses affects the development of the bacterial microbiome.

Breast milk is known to have a protective effect against human pathogenic viruses and is associated with decreased infant mortality in developed and developing countries (Liang et al., 2020). Many studies, however, have identified enteric pathogenic viruses being asymptomatically carried in otherwise healthy children (McCann et al., 2018; Liang et al., 2020; Kapusinsky et al., 2012; Tan et al., 2020; Motogosi et al., 2020). Comparable to what was described by Tan et al. (2020) in Bangladesh, we found that picornaviruses dominated the infant gut human-host eukaryotic virome, particularly parechoviruses and enteroviruses, suggesting that picornaviruses are common virome colonizers regardless of geographic region. Infants also asymptomatically bore noroviruses, caliciviruses, astroviruses and rotaviruses. Asymptomatic carriage of these viruses has been reported by Taboada et al. (2020) in a cohort of three infants in semi-rural Mexico, which may share cultural and nutritional similarities to the cohort described in this study. While the association with the detection of virus and the diagnosis of illness is beyond the scope of this paper and will be the focus in future reports, 48/52 (92.3%) of the infants reported illness over the three years. In the current study, all Infants were asymptomatic at the time of sample collection, indicating that our results were likely not skewed by an active infection, including the one infant with an overabundance of parechovirus reads at the B7 timepoint (Figure 4). In comparison to infants, we observed that the maternal virome is stable in composition for over a year, during the third trimester and 1-year post-partum. Changes to the maternal gut microbial community have only been documented in the bacterial microbiome, reporting an increase in proteobacteria and actinobacteria and reduced richness from the first to third trimester (Koren et al., 2012).

We report that both the prokaryotic and eukaryotic components of the virome are distinct between mother and child in early life. Viral abundance in infants increased over time, as reported in other studies (Yatsunenko et al., 2012; Maqsood et al., 2019). While the infant prokaryotic virome became adult-like in composition by the third year, the infant eukaryotic virome remained distinct, consisting primarily of picornaviruses and anelloviruses, as compared to retroviruses and herpesviruses in the mother. Maqsood et al. (2019) determined that the neonatal virome is only 15% related to the mother’s virome, and concluded that colonization most likely occurs through other environments like breast milk, skin, and surfaces. Here we also found that the diversity of the infant virome was not primarily driven by exposure to the mother but rather, is more likely to be determined by exposure to other household members and/or the environment. Future studies are needed to determine how the presence / absence or persistence of certain pathogenic viruses contributes to disease later in life.

Our findings are limited by indirect comparisons of the infant to the adult virome in the third year, a lack of early pregnancy samples in the mothers, a lack of analysis of male guardians/parents and limited racial diversity in our population. However, the multiple timepoints collected in both infants and mothers allow us to observe meaningful trends in the virome. Future investigations of the STORK cohort will investigate the mechanics of bacterial and viral interactions, infection and the virome, and the relationship between the virome and weight gain and immune development. In conclusion, we determined the infant virome changes dramatically from the first month to the third year; however, the infant virome at age three years still retains a distinct composition and diversity from their mothers’.

## Data Availability

Preprocessed reads with human sequences removed by high-sensitivity local alignment to the human genome (GRChg38/hg38 build) using Bowtie2(Langmead and Salzberg, 2012) have been deposited into the NCBI Sequence Read Archive (SRA) (accession numbers pending).

## Contributions

J.P. and C.Y.C. conceived and designed the experiments. A.C.G., C.L. T.H., and A.S.-G. performed the experiments. W.A.W., S.F., Y.S., V.S., R.E.L., J.P., and C.Y.C. analyzed data. R.E.L., J.P., and C.Y.C. contributed reagents/materials/analysis tools. A.C.G., J.P., and C.Y.C. wrote the paper. All authors reviewed the manuscript and agreed to its contents.

## Contributions

The authors declare that there are no competing interests.

## Acknowledgments

We thank the staff at the UCSF Center for Advanced Technology for sequencing transcriptome libraries using the Illumina NovaSeq 6000.The NIH and the Max Planck Society funded this study.

---

**Supplemental Figure 1.**
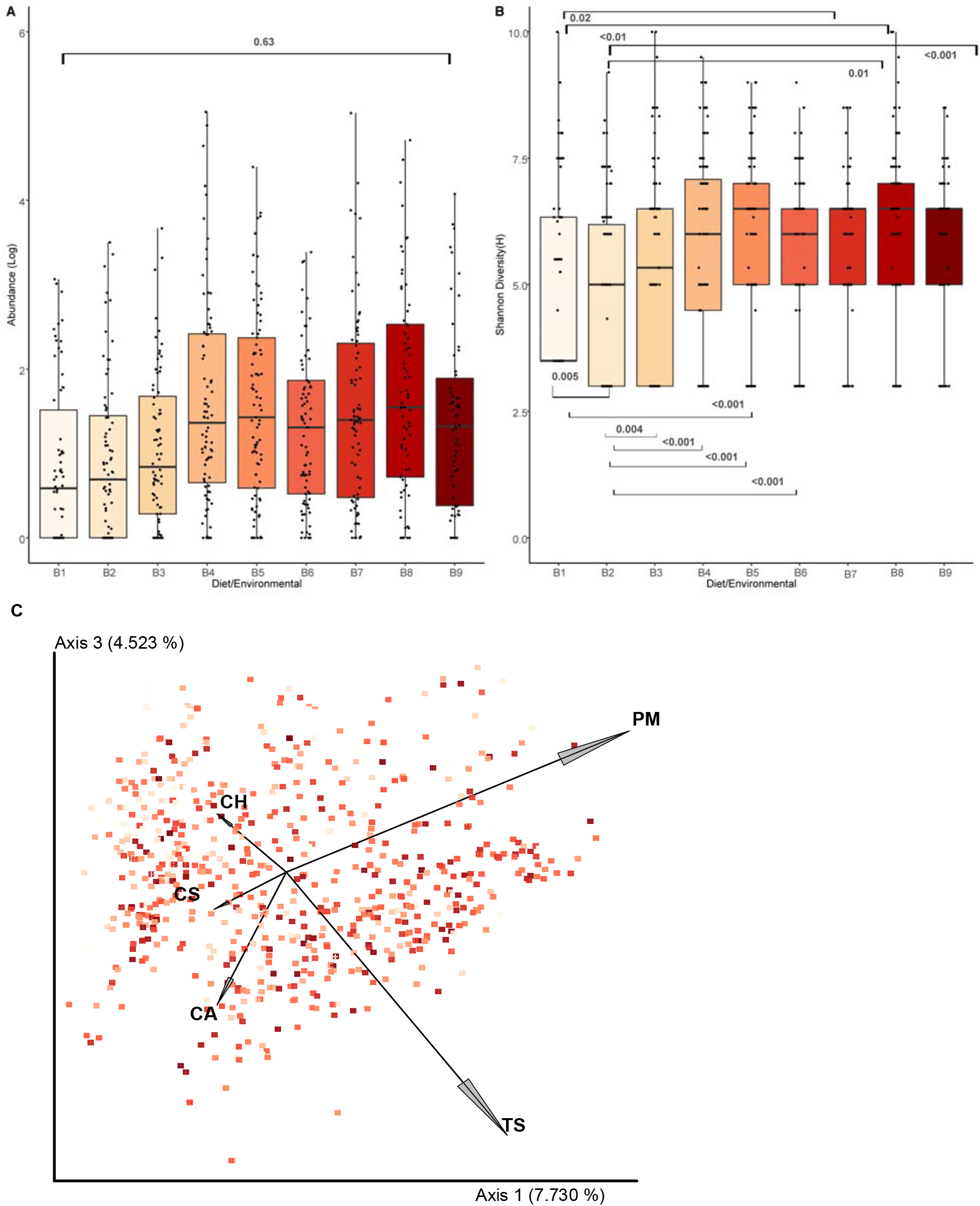
Normalized abundance, alpha and beta diversity of dietary/environmental viruses in infants. Box and whiskers plot of **(A)** dietary/environmental virus abundance and **(B)** Shannon diversity**. (C)** PCA plot of infant samples from dietary/environmental viral counts showing Jaccard distances. Samples are colorized with an age gradient (light to dark red representing age in days of the infants, range of 22 to 1166). The top 6 taxa driving clustering are shown as a biplot. Abbreviations: PM: pepper mild mottle virus, TS: tropical soda apple mosaic virus, TM: tomato mosaic virus, CS: chicken stool-associated gemycircularvirus, CA: chicken anemia virus.

**Supplemental Figure 2.**
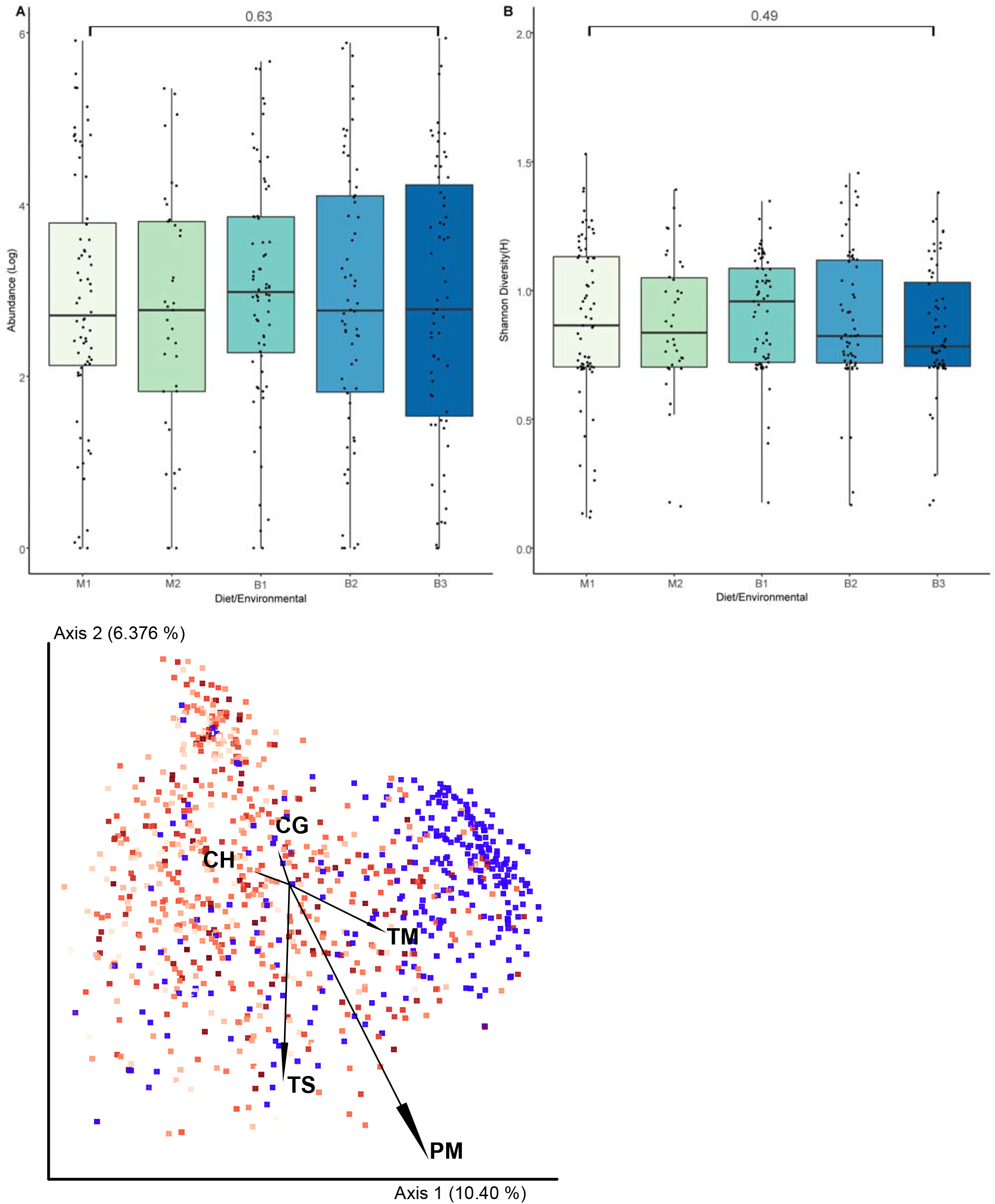
Normalized abundance, alpha and beta diversity of dietary/environmental viruses in mothers. Box and whiskers plot of **(A)** dietary / environmental virus abundance and **(B)** Shannon diversity. **(C)** PCA plot of infant samples from dietary/environmental viral counts showing Jaccard distances. Samples are colorized with an age gradient (light to dark red representing age in days of the infants, range of 22 to 1166 and mothers, colored in blue. The top 5 taxa driving clustering are shown as a biplot. Abbreviations: PM: pepper mild mottle virus, TS: tropical soda apple mosaic virus, TM: tomato mosaic virus, CH: cyprinid herpesvirus 3, CG: cucumber green mottle mosaic virus.

**Supplemental Figure 3.**
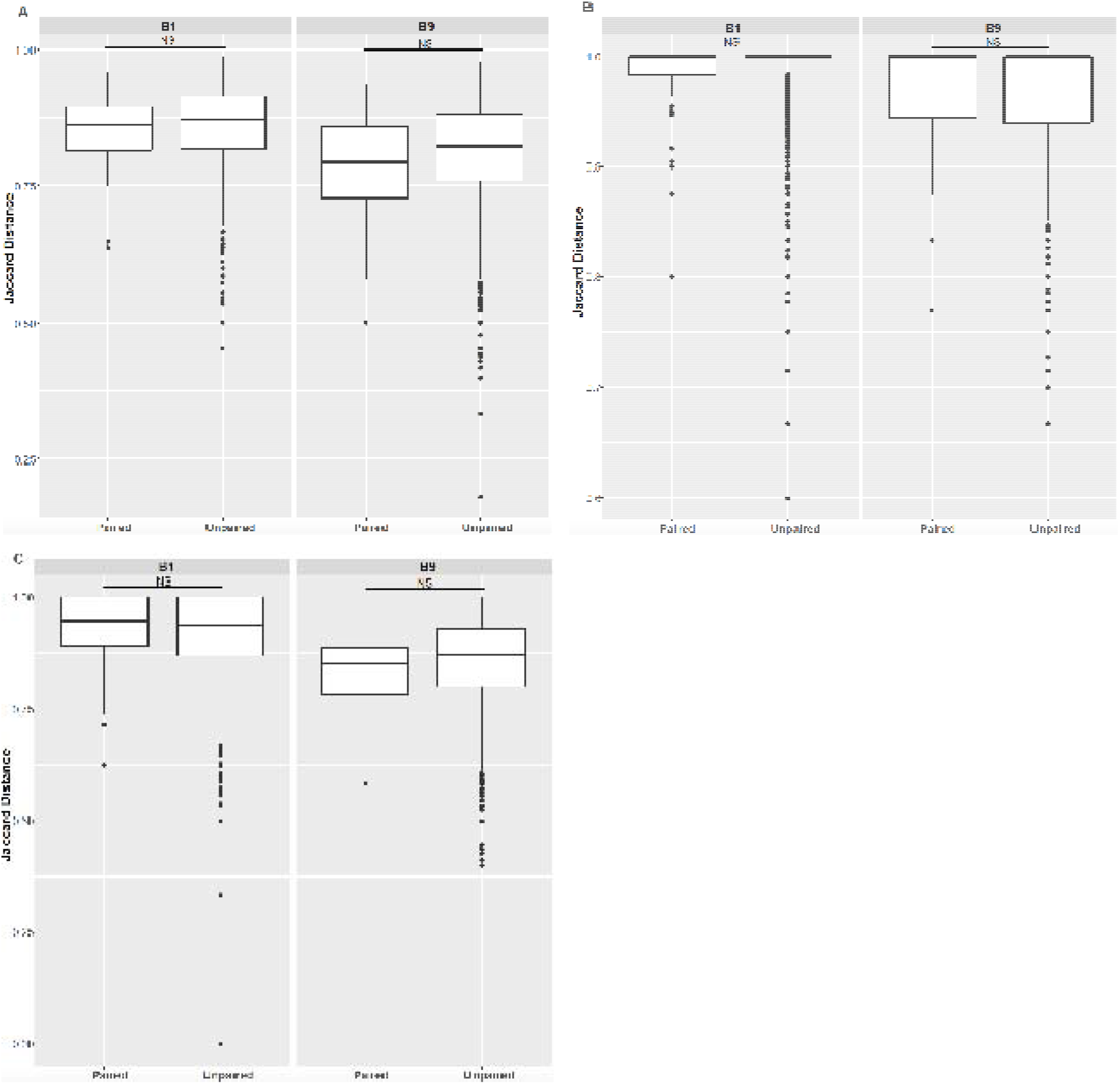
Jaccard distances between infant / mother pairs and unpaired infants and mothers. Jaccard distances at time points B1 and B9 are shown for (**A**) prokaryotic viruses, (**B**) eukaryotic virus, and (**C**) dietary/environmental viruses.

